# Oxygen plasma focused ion beam scanning electron microscopy for biological samples

**DOI:** 10.1101/457820

**Authors:** Gorelick Sergey, Korneev Denis, Handley Ava, Gervinskas Gediminas, Oorschot Viola, Kaluza Owen L., Law Ruby H.P., Bryan Moira O’, Pocock Roger, Whisstock James C., de Marco Alex

**Affiliations:** ARC Centre of Excellence in Advanced Molecular Imaging; Biomedicine Discovery Institute, Department of Biochemistry and Molecular Biology, Monash University, Clayton Australia; School of Biological Sciences, Monash University, Clayton Australia; Biomedicine Discovery Institute, Department of Anatomy and Developmental Biology, Monash University, Clayton Australia; Clive and Vera Ramaciotti Centre for Cryo-Electron Microscopy, Monash University, Clayton Australia; Monash eResearch Centre, Monash University, Melbourne Australia; EMBL Australia, Monash University, Clayton Australia; University of Warwick, Coventry CV4 7AL, United Kingdom

**Keywords:** O-PFIB, oxygen plasma FIB, FIB/SEM tomography, Neurobiology, Correlative Microscopy, imaging, cell biology

## Abstract

Over the past decade, gallium Focused Ion Beam-Scanning Electron Microscopy (FIB-SEM) has been established as a key technology for cellular tomography. The utility of this approach, however, is severely limited both by throughput and the limited selection of compatible sample preparation protocols. Here, we address these limitations and present oxygen plasma FIB (O-PFIB) as a new and versatile tool for cellular FIB-SEM tomography. Oxygen displays superior resin compatibility to other ion beams and produces curtain-free surfaces with minimal polishing. Our novel approach permits more flexible sample preparation and 30% faster data collection when compared to using gallium ion sources. We demonstrate this alternative FIB is applicable to a variety of embedding procedures and biological samples including brain tissue and whole organisms. Finally, we demonstrate the use of O-PFIB to produce targeted FIB-SEM tomograms through fiducial free *en-block* correlative light and electron microscopy.

## Main

Electron microscopy (EM) is one of the most commonly used approaches for imaging the cellular environment at the nanometre scale. The main limitation of cellular EM is that acquiring statistically relevant datasets, large areas or sparse and rare events is extremely time-consuming – a single dataset easily taking more than one week of imaging time. For example, most high-resolution three-dimensional (3D) cellular imaging to date has been performed through transmission electron tomography [1], which requires sample sectioning. Here, if the region of interest is larger than 200-300 nm (700 nm if using Scanning Transmission Electron Tomography [2]), the procedure requires laborious identification and imaging of the same region across multiple sections [3].

With a few exceptions [4], the best performing imaging approaches for large volumes are based on the use of Scanning Electron Microscopy (SEM) either in conjunction with serial sectioning (e.g. 3View™, VolumeScope™, ATUMtome™) [5, 6] or with Focused Ion Beam (FIB) milling [7]. Of these techniques, only the combination/union of FIB and SEM techniques allows imaging at sub-10 nm resolution in 3D [8, 9].

The combination of serial block-face sectioning and imaging allows fast and precise removal of material from the block face and imaging of the newly exposed face. The limitations here are linked to the stiffness of the resins that can be used (typically epoxybased such as Epon™ or Durcupan™), the thickness of the sections that can be reproducibly achieved (30-50 nm) [10] and negative surface charging that greatly influences image quality.

FIB-SEM tomography utilizes focused electron and ion beams that are both pointing at the same location on the sample but from different angles [11] (figure S1). In this case, the block face around the region of interest (ROI) is milled away using the focused ion beam (instead of a diamond knife) and the newly exposed face is imaged by means of the electron beam (figure S1). This method has the major advantage that it is not limited to a particular geometry of the sample, and the imaged face can be prepared at any angle relative to a preexisting sample surface [8, 12]. The flexibility of FIB-SEM tomography also includes slice thickness, that can range from 1 nm to multiple microns (optimally between 5 nm and 100 nm). The major limitations of this technique are linked to the types of resin that can be successfully employed when embedding biological material and to the throughput, which with the exception of few reported cases [13], is generally limited to ~5 h / μm upon the acquisition of an area of 30×30 μm with a voxel size of ~5 nm.

The most common FIB system, used throughout the materials science community, is based on a liquid metal source (typically gallium) capable of providing a fine and well-defined beam (with resolution down to 2-5 nm) [14]. Although this technology has been successfully adapted to selected cell biology studies, as shown by the published volume sizes [10], it does not cope well with the removal of sample volumes which are larger than 50’000 μm^3^. Furthermore, the interaction between the slicing metallic ion beam and organic samples has never been optimized.

FIB systems based on inductively coupled plasma (ICP) sources [15, 16] (plasma FIBs, PFIBs) allow a wide range of ion beam species (e.g. noble Ar and Xe, O and N) which eliminates the effect of metal contamination on the milled surface due to implantation [17]. ICP sources are capable of delivering orders of magnitude higher currents (up to 2.5 μA versus 65 nA on traditional liquid metal Ga-FIB) and still maintain a micrometre spot-size due to the larger source size, which results in lower visible spherical aberration. There have been a few reports of PFIB successfully used to enhance the throughput of FIB-SEM tomography in material sciences [18], however, these ICP beams have not yet been extensively applied to imaging of biological samples.

Here, we present the use of an O plasma-FIB (O-PFIB) as an alternative and optimized beam to be used when milling carbon-based samples [19]. Our results show that O-PFIB has superior compatibility with all resins commonly used in biology, including acrylic-based resin. We further show that as a result of increased resin compatibility, existing protocols for Correlative Light and Electron Microscopy (CLEM) [20] that were restricted to TEM can be used to target regions in a sample which have been marked with commonly used fluorescent tags such as GFP. In addition, we show that the use of non-metal ion beam sources enhances imaging speed due to the reduction of background noise.

## Results and Discussion

FIB-SEM tomography is a highly repetitive process that comprises a preparatory phase and an imaging phase of the ROI (figure S1) [7]. The preparatory phase includes FIB milling, area alignment (optional) and focussing (optional). In a typical cellular imaging setup, where the needed pixel-size is approximately 5 nm, one would expose an area 17-20 μm wide and ~15 μm deep; while in a tissue context the area for the same required pixel size would be ~35-40 μm wide and 30 μm deep. All the above-mentioned values are only indicative and can become 2-4 times larger, but in this work, we focussed on optimising the most commonly imaged volume sizes.

Since the milling step is preparatory and does not provide data per se, it has been shown that an optimal milling procedure should not require more than 20% of the SEM imaging time [13]. Thus, in the situation where an area of 35×30 μm^2^ is imaged with a pixel size of 5.6 nm and the imaging dwell time is 1 μs the acquisition will require ~25 sec. Here the milling procedure should last no longer than 5 sec. Looking at the most commonly used milling conditions (on Zeiss and ThermoFisher Ga based ion columns) the used currents are between 0.7-27 nA [7, 13, 21]. The higher currents in this range work efficiently in geometries where the block face is milled from glancing angle, whereas if the ROI is located within a trench on the block face (figure S1), in order to limit the damage and the surface roughness (known as curtaining) lower currents are required.

In this work, we used ThermoFisher Helios G4 instruments (Ga FIB and PFIB) to investigate if there is an optimised ion beam chemistry to be used on resin embedded biological samples. Using a prototype PFIB capable of switching the plasma source (Ar, Xe and O) we were able to efficiently expose the ROI with ion currents as high 60 nA (figure 1) and on Ga-FIB using current as high as 20 nA. These experiments permitted cutting with a resolution of 10 nm if the geometry of the sample permitted glancing access to the block face. We tested and compared the behaviour upon exposure of four commonly used resins (Durcupan™, Epon™, HM20™, and LR-White™) to different ion species and currents. Collectively, these data reveal that Xe and Ga only perform well on Epon and Durcupan regardless of the current used. Instead, the softer resins suffer from curtaining even at currents as low as 4 nA on Xe (2.4 nA on Ga). At 4 nA, Ar performs well on Durcupan, Epon and HM20 whilst at 60 nA Ar only performs well on Durcupan. In contrast, O plasma was compatible with all resins tested and was curtain-free were, regardless of the milling current (up to 60 nA, see figure 1).

**Figure 1:**
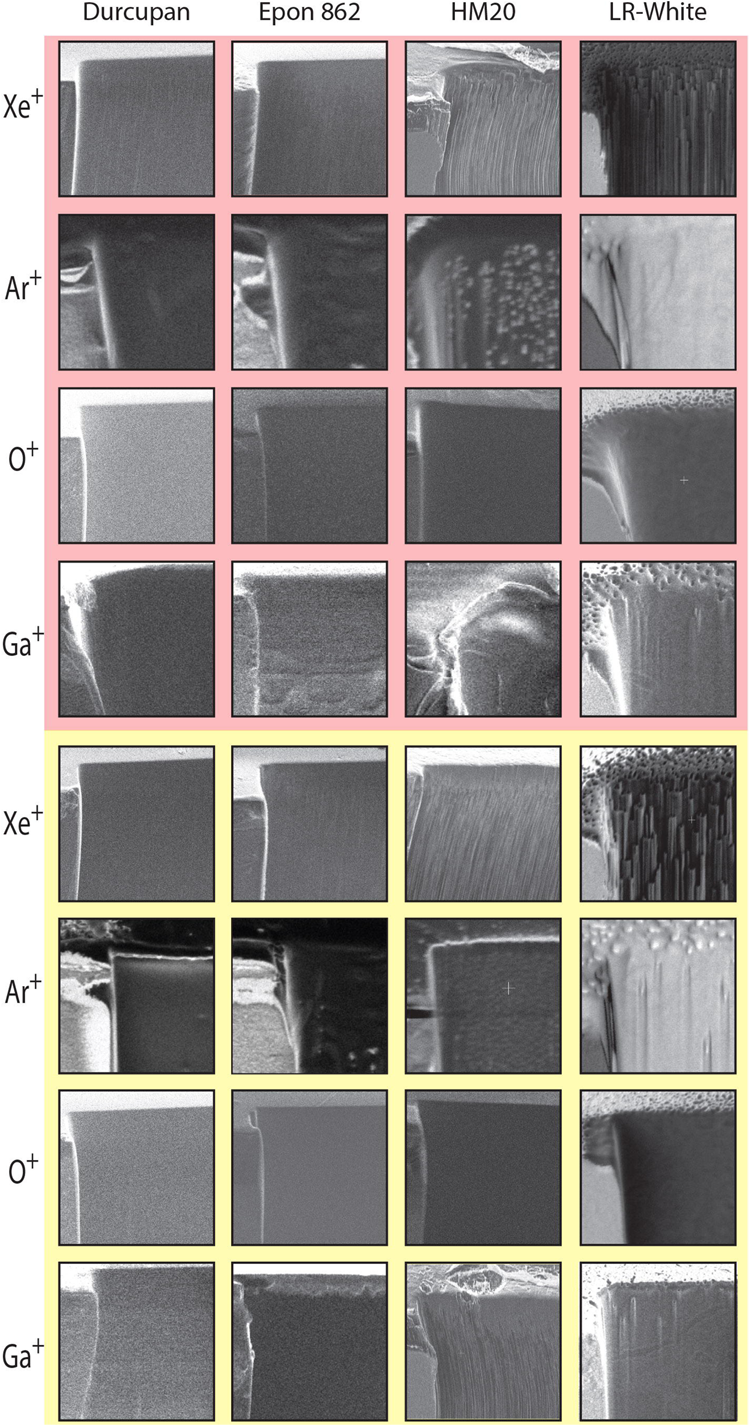
Beam – resin compatibility. tested on 3 ICP and 1 metal ion sources. Displayed in the red panel is a cut performed at the maximum current that can be used for face preparation when exposing the region of interest (60 nA for ICP sources and 20 nA for Ga). In the yellow panel the best-performing current (which keeps the milling time for a 30 × 30 × 0.01 pm face below 10 s). 4 nA for Xe, 15 nA for Ar and O and 9 nA for Ga. Notably, with the exception of Durcupan, which appears to be compatible with every beam, O appears to have the best compatibility.

The majority of the time required when milling a cross-section with a FIB is not devoted to the removal of material but rather to the surface finish (figure S3). Our results show that milling a resin with different ion species results in significant differences in the surface finish. The fact that the surfaces produced with O beam are free from curtains already after 3-5 seconds, it makes possible to obtain a surface suitable for SEM imaging in significantly shorter time (figure 1, S3).

We next investigated the relationship between ion current and milling efficiency. To achieve this, we first measured the ion beam spot size yielded by both Ga FIB and PFIB at different currents (figure S2A). As expected, the spot size of the Ga beam rapidly deteriorates with the increase of the current because of the spherical aberration. In contrast, the Ar, Xe and O spot size is source-limited at currents lower than 4nA, but it is remarkably better focused at currents higher than 12 nA (figure S2). We then simulated and measured the sputter rate of all used ions at 30 keV and incident at an angle normal to the surface on Epon 812. All simulations were performed using SRIM [22], while direct measurements were acquired by exposing a 5×5 or 20×20 μm squares to ion fluencies from 10^4^ to 10^6^ nC at different current densities and by measuring the removed volume (figure S2 B). Unexpectedly, we found that the simulated and measured sputter rates were comparable for all ions with the exception of O. Here, the sputter yield is increased by a factor of 3 compared to the simulation based on the atomic number. Since oxygen is a reactive gas, we hypothesise that its interaction with C or C-N based substrate may lead to the formation of volatile compounds in a process similar to that of reactive ion etching [23]. Such an event would invalidate the commonly used sputtering calculations and a chemical etching component should be taken into consideration. A second point to consider is that the ICP plasma source yields a substantial secondary spot that is composed of ionized molecular oxygen (O_2_+). Since both spots coincide at the focus the sputtering rate is partly modified.

Given the fact that O is the best performing beam, we tested the utility of the O beam on mouse brain tissue embedded in different resins (figure 2). For HM20, uranyl acetate staining was used instead of Osmium (because of the incompatibility of this resin with OsO4 staining). Comparisons of the FIB-SEM tomography slices reveal that excellent quality images can be obtained no matter the resin type or staining procedure. For example, all expected organelles such as the Golgi are visible, and features such as nuclear pores (arrowed; panel 2) can clearly be distinguished.

The majority of the time required when milling a cross-section with a FIB is not devoted to the removal of material but rather to the surface finish (figure S3). Our results show that milling a resin with different ion species results in significant differences in the surface finish. The fact that the surfaces produced with O beam are free from curtains already after 3-5 seconds, it makes possible to obtain a surface suitable for SEM imaging in significantly shorter time (figure 1, S3).

We next investigated the relationship between ion current and milling efficiency. To achieve this, we first measured the ion beam spot size yielded by both Ga FIB and PFIB at different currents (figure S2A). As expected, the spot size of the Ga beam rapidly deteriorates with the increase of the current because of the spherical aberration. In contrast, the Ar, Xe and O spot size is source-limited at currents lower than 4nA, but it is remarkably better focused at currents higher than 12 nA (figure S2). We then simulated and measured the sputter rate of all used ions at 30 keV and incident at an angle normal to the surface on Epon 812. All simulations were performed using SRIM [22], while direct measurements were acquired by exposing a 5×5 or 20×20 μm squares to ion fluencies from 10^4^ to 10^6^ nC at different current densities and by measuring the removed volume (figure S2 B). Unexpectedly, we found that the simulated and measured sputter rates were comparable for all ions with the exception of O. Here, the sputter yield is increased by a factor of 3 compared to the simulation based on the atomic number. Since oxygen is a reactive gas, we hypothesise that its interaction with C or C-N based substrate may lead to the formation of volatile compounds in a process similar to that of reactive ion etching [23]. Such an event would invalidate the commonly used sputtering calculations and a chemical etching component should be taken into consideration. A second point to consider is that the ICP plasma source yields a substantial secondary spot that is composed of ionized molecular oxygen (O_2_+). Since both spots coincide at the focus the sputtering rate is partly modified.

Given the fact that O is the best performing beam, we tested the utility of the O beam on mouse brain tissue embedded in different resins (figure 2). For HM20, uranyl acetate staining was used instead of Osmium (because of the incompatibility of this resin with OsO4 staining). Comparisons of the FIB-SEM tomography slices reveal that excellent quality images can be obtained no matter the resin type or staining procedure. For example, all expected organelles such as the Golgi are visible, and features such as nuclear pores (arrowed; panel 2) can clearly be distinguished.

The above results demonstrate the ability to use oxygen as a versatile ion beam on biological samples providing an increased compatibility with sample preparation protocols. Next, we investigated the speed of image acquisition and compared the quality of images obtainable after milling with other ion species. In material sciences, Ga implantation is a known phenomenon that results from the use of Ga-FIB [17, 24]. In biological samples, the effect of Ga implantation is not obvious and therefore has always been disregarded. Here we measured a clear improvement in the backscatter contrast on OsO_4_ stained brain sample embedded in Epon when milling with non-metallic ion beams (figure 3). Our results show that the contrast between the stained and unstained regions is clearly decreased in the images acquired after milling with Ga when compared to the contrast from samples milled with the plasma beams (figure 3). This effect was quantified as local variance in the grey values of the backscatter image (figure S3 B, C). In order to obtain comparable quality images, a dwell time of 1 μs is required for samples produced using the Ga FIB versus ~700 ns dwell time for samples milled using non-metallic ion beams. Thus, using of non-metallic ion beams enables to reduce the image acquisition time by ~30%, which predictably results in ~30% larger volumes imaged in an allocated session (figure S3).

In addition to the direct analysis of image noise, we verified the detrimental effect of Ga implantation on the overall backscatter electron image by performing Monte Carlo simulations [25]. Here, statistics of 2 keV electrons backscattered from Epon with varying concentrations of Os, Ga, as well as their mixtures were analysed to estimate the probability of an electron to backscatter and contribute to the pixel signal. The implanted Ga atoms, although lighter than the Os stain, are nevertheless considerably heavier than the primary atomic constituents of the Epon host resin (C, O and H). Thus, Ga generates a significant contribution to the backscatter signal above the uniform background signal emanating from the resin. We found that Ga implantation produces >25% of the signal associated with a similar concentration of Os atoms (figure S3). Hence, the effect of Ga-dilution of Os in Epon is to decrease the observed backscatter image contrast.

The versatility that O plasma beams bring toward various resins opens the possibility of developing FIB-SEM tomography for protocols that were previously restricted to TEM (transmission electron microscopy) or array tomography. To demonstrate this capability, we tested the use of O beam in Correlative Light and Electron Microscopy (CLEM) experiments. We adapted the approaches, described in Kukulski et al. [26] and Peddie et al. [27], where samples are subjected to high-pressure freezing, freeze substitution, staining with uranyl acetate and infiltration with HM20 resin. This preparation does not make use of OsO4 until after the light microscopy has been performed because it would shield fluorescence, and as a result, the cellular membranes are not visible under TEM. Accordingly, after fluorescence imaging and in preparation for EM imaging, the sections are post-stained with OsO4 and or lead citrate [27].

As discussed earlier, Ga FIB-SEM cannot be used to efficiently mill samples embedded in HM20. In contrast, we demonstrated that uranyl acetate stained samples embedded in HM20 can be efficiently milled using the O-PFIB and imaged under the SEM (figure 2). Accordingly, we sought to demonstrate the use of O-PFIB for targeted imaging using CLEM using an approach similar to the one described by Hohn et al. [28], but resulting free from curtains and milling artefacts. We performed *en-block* confocal imaging of a region of high pressure frozen and freeze substituted HM20 embedded and uranyl acetate stained *Caenorhabditis elegans.* From the confocal stack, it was possible to identify individual cells and the resolution was good enough to identify sensory dendrites in the worm head {Juozaityte, 2017 #4}(figure 4E). Most importantly, it was possible to estimate the depth of the region of interest from the block surface, which provided the knowledge about how much milling was required to expose the target region (in this case 46 μm). The spatial resolution was estimated to be ~300 nm in XY and ~600 nm along Z. Considering a target imaging field in the FIB-SEM of ~30 μm, this provided an excellent approach for targeted FIB-SEM tomography. Accordingly, we transferred the sample into the O-PFIB and targeted the acquisition of a tomogram of a 30 × 30 × 15 μm portion of the intestinal region of an entire L1 larva. O-PFIB-SEM data collection took 16 hours, and the resulting tomogram was produced with ~10 x ×10 x ×15 nm resolution and it is curtain free (figure 4F, video S1). The possibility to acquire targeted datasets once more can result into a significant increase of the throughput.

In conclusion, we demonstrate that in contrast to all ions tested, oxygen represents a superior beam for milling the most currently used resins in the life sciences. In addition, the O-PFIB-SEM provides superior results in terms of data acquisition efficiency and image quality. Finally, we show that O-PFIB can be used to readily yield FIB-SEM tomograms in the context of CLEM experiments – opening up previously unavailable possibilities in this field such as targeted high-resolution imaging in large samples.

## Methods

### Resin embedding

Animal tissue was humanely collected in accordance with the National Health and Medical Research (NHMRC) guidelines and as approved by the Monash University Biological Sciences Animal Ethics Committee (BSCI/2017/46).

Mouse brain tissue was fixed in either 2.5% glutaraldehyde, 2% paraformaldehyde in 0.1M sodium cacodylate buffer or fixed in 2%PFA 0.2% GA in 0.1M Phem. After initial fixation tissue was post-fixed with 1% OsO4, 1.5% K3Fe(III)(CN)6. Dehydration was done at room temperature by increasing the concentration of ethanol. Sample embedding in the resin was done at room temperature through a series of steps with increasing the resin in ethanol: 25% then 50% and 75% followed by 2x 100% resin. Blocks were polymerized for 48-56 h at 60 °C. Epon 812 (EMS14120), Durcupan (Sigma-Aldrich 44610) or LR-White (EMS 14380).

### Lowicryl HM20 embedding

Mouse brain tissue was fixed in 4% paraformaldehyde in 0.1M PHEM buffer, infused in 2.3M sucrose and frozen in Liquid Nitrogen. Automatic Freeze Substitution (Leica EM AFS2) was done in 0.1% Uranyl Acetate in acetone (−90°C till −50°C), followed by intensively rinsing in acetone. Sample embedding in HM20 was done at −50°C through a series of steps of HM20 in acetone: 25% then 50% and 75% followed by 5x 100% Lowicryl HM20 (EMS 14340). UV polymerization was performed at -50°C for 5 days.

### FIB-SEM tomography

FIB-SEM tomography experiments were performed using a ThermoFisher Helios UX G4 or a prototype ThermoFisher Multigas Helios™ UX G4 PFIB capable of delivering switchable plasma sources. In all cases, milling was performed at 30 keV and ion currents in use were between 10 pA and 60 nA.

All trenches and initial block face polishing were milled using a standard cleaning crosssection pattern. Currents were measured prior to the experiments using a Faraday cup to make sure the nominal current would not differ more than 10% from the measured beam current. Imaging was performed to detect Backscattered electrons through the in-column detector or the Through Lens Detector (TLD) with the electrostatic immersion field on. Dwell time for imaging was kept between 200 ns and 3μs. Pt deposition for the protection layer was performed through the GIS (gas injection system) either using the ion beam induced deposition at 30 keV and 60 nA or the electron beam induced deposition at 2 keV at 25 nA. Samples were glued on a standard SEM stub using Epoxy glue and subsequently coated with 5 nm of Ir to minimize charging effects.

Automated FIB-SEM tomography routines from ThermoFisher AutoSlice & View™ or iFAST™ were used to collect stacks.

### Image quality/noise comparison

All data were acquired on the same SEM column (FEI Elstar™ G4) at 4 mm working distance, 2 keV accelerating voltage, 400 pA electron beam current and dwell time between 300 ns and 1 μs. The test sample was mouse brain stained with OsO_4_ and embedded in Epon. The images acquired after milling with Xe, O, and Ar were prepared on the same instrument on the same day, while the ones milled with Ga were at first measured on a separate instrument, but in the second instance, the images were reacquired on the same microscope as the first three beams. Ga implantation was confirmed using EDX. Although Ga presence could be detected, no quantification was possible because the resin could not sustain the required electron dose.

### Image processing

Stacks were filtered and aligned using custom algorithms written in Python (http://www.python.org) following the approach described in Saalfeld et al. [29]. Visualization and image analyses were conducted using ThermoFisher Avizo™, Matlab™ (MathWorks) and Fiji [30]. Noise analyses were performed through custom algorithms written in Python and Matlab. All images were mean-normalized prior to noise analysis. Figures were prepared using Matlab and annotated using Adobe Illustrator™.

### Correlative light and electron microscopy

For the purpose of targeting the FIB-SEM tomography acquisition to a specific site, we used fluorescently labelled *C. elegans (RJP3088 Pets-5::mCherry + Pelt-2::GFP [31]).* Worms were maintained at 20 °C on NGM plates seeded with *Escherichia coli* 0P50. L1 larvae were isolated as described in Porta-de-la-Riva et al. [32]. Worms were suspended in a solution of 20% BSA in M9 and pipetted on a Leica EM-PACT2 carrier. We proceeded with freeze substitution and HM20 embedding as described in Kukulski et al. [26]. After polymerization, all blocks were hand-trimmed to ensure a certain degree of asymmetry in the shape of the block face. Confocal imaging was performed on an upright Leica SP8 scanning confocal, using a 20× / 1 NA water immersion objective.

**Figure 2:**
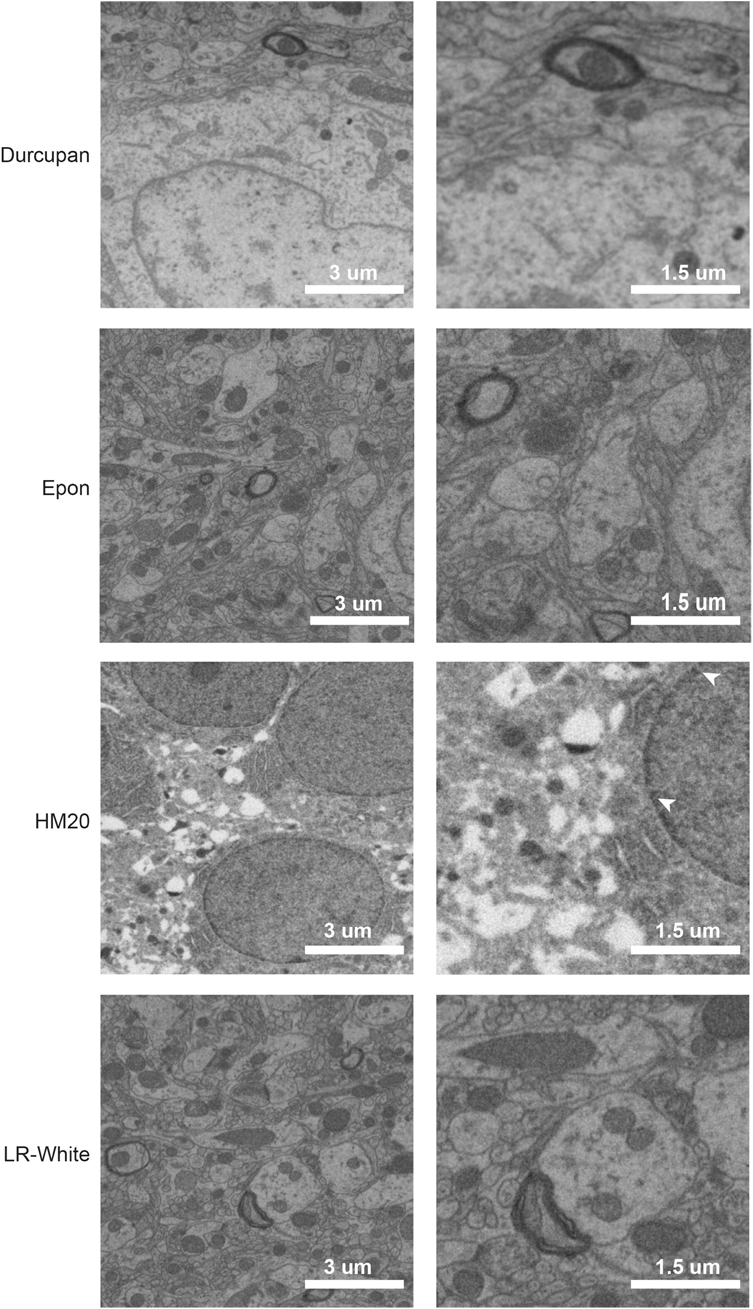
FIB-SEM tomography of mouse brain embedded in different resins using oxygen plasma to expose the imaged surface. All samples were stained with OsO4 with the exception for the one embedded in HM20. Notably, all images were collected with the same imaging conditions (current, WD, dwell, orientation) but the contrast appears to be significantly different.

The above results demonstrate the ability to use oxygen as a versatile ion beam on biological samples providing an increased compatibility with sample preparation protocols. Next, we investigated the speed of image acquisition and compared the quality of images obtainable after milling with other ion species. In material sciences, Ga implantation is a known phenomenon that results from the use of Ga-FIB [17, 24]. In biological samples, the effect of Ga implantation is not obvious and therefore has always been disregarded. Here we measured a clear improvement in the backscatter contrast on OsO_4_ stained brain sample embedded in Epon when milling with non-metallic ion beams (figure 3). Our results show that the contrast between the stained and unstained regions is clearly decreased in the images acquired after milling with Ga when compared to the contrast from samples milled with the plasma beams (figure 3). This effect was quantified as local variance in the grey values of the backscatter image (figure S3 B, C). In order to obtain comparable quality images, a dwell time of 1 ps is required for samples produced using the Ga FIB versus ~700 ns dwell time for samples milled using non-metallic ion beams. Thus, using of non-metallic ion beams enables to reduce the image acquisition time by ~30%, which predictably results in ~30% larger volumes imaged in an allocated session (figure S3).

**Figure 3:**
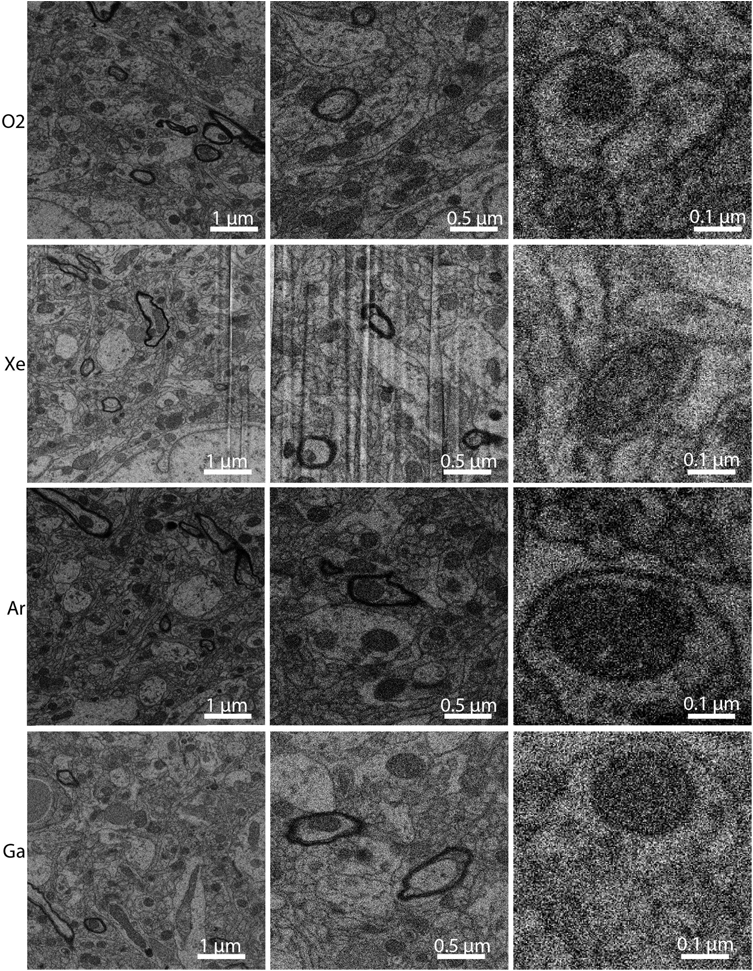
The effect of metal implantation on backscatter contrast in heavy metal stained samples embedded in Epon. Notably, when milling with non-metal beams the contrast appears increased. The heavy curtaining displayed by Xe makes this beam not optimal for imaging on this resin.

In addition to the direct analysis of image noise, we verified the detrimental effect of Ga implantation on the overall backscatter electron image by performing Monte Carlo simulations [25]. Here, statistics of 2 keV electrons backscattered from Epon with varying concentrations of Os, Ga, as well as their mixtures were analysed to estimate the probability of an electron to backscatter and contribute to the pixel signal. The implanted Ga atoms, although lighter than the Os stain, are nevertheless considerably heavier than the primary atomic constituents of the Epon host resin (C, O and H). Thus, Ga generates a significant contribution to the backscatter signal above the uniform background signal emanating from the resin. We found that Ga implantation produces >25% of the signal associated with a similar concentration of Os atoms (figure S3). Hence, the effect of Ga-dilution of Os in Epon is to decrease the observed backscatter image contrast.

The versatility that O plasma beams bring toward various resins opens the possibility of developing FIB-SEM tomography for protocols that were previously restricted to TEM (transmission electron microscopy) or array tomography. To demonstrate this capability, we tested the use of O beam in Correlative Light and Electron Microscopy (CLEM) experiments. We adapted the approaches, described in Kukulski et al. [26] and Peddie et al. [27], where samples are subjected to high-pressure freezing, freeze substitution, staining with uranyl acetate and infiltration with HM20 resin. This preparation does not make use of OsO4 until after the light microscopy has been performed because it would shield fluorescence, and as a result, the cellular membranes are not visible under TEM. Accordingly, after fluorescence imaging and in preparation for EM imaging, the sections are post-stained with OsO4 and or lead citrate [27].

As discussed earlier, Ga FIB-SEM cannot be used to efficiently mill samples embedded in HM20. In contrast, we demonstrated that uranyl acetate stained samples embedded in HM20 can be efficiently milled using the O-PFIB and imaged under the SEM (figure 2). Accordingly, we sought to demonstrate the use of O-PFIB for targeted imaging using CLEM using an approach similar to the one described by Hohn et al. [28], but resulting free from curtains and milling artefacts. We performed *en-block* confocal imaging of a region of high pressure frozen and freeze substituted HM20 embedded and uranyl acetate stained *Caenorhabditis elegans.* From the confocal stack, it was possible to identify individual cells and the resolution was good enough to identify sensory dendrites in the worm head {Juozaityte, 2017 #4}(figure 4E). Most importantly, it was possible to estimate the depth of the region of interest from the block surface, which provided the knowledge about how much milling was required to expose the target region (in this case 46 μm). The spatial resolution was estimated to be ~300 nm in XY and ~600 nm along Z. Considering a target imaging field in the FIB-SEM of ~30 μm, this provided an excellent approach for targeted FIB-SEM tomography. Accordingly, we transferred the sample into the O-PFIB and targeted the acquisition of a tomogram of a 30 × 30 × 15 μm portion of the intestinal region of an entire L1 larva. O-PFIB-SEM data collection took 16 hours, and the resulting tomogram was produced with ~10 x ×10 x ×15 nm resolution and it is curtain free (figure 4F, video S1). The possibility to acquire targeted datasets once more can result into a significant increase of the throughput.

**Figure 4:**
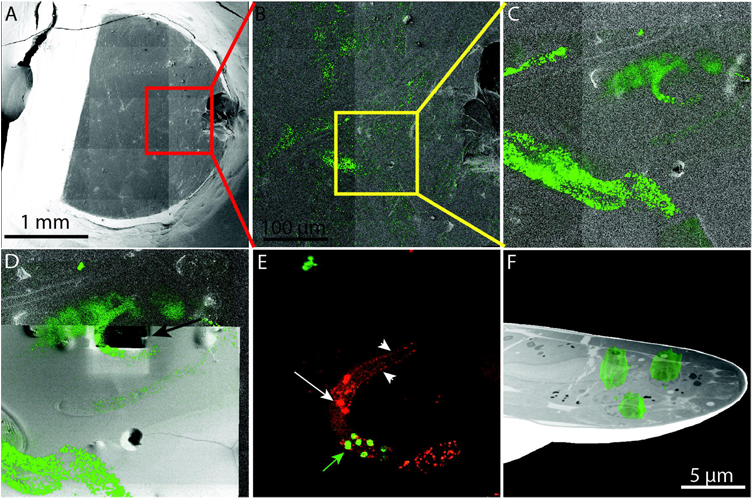
*En-block* Correlative Light and Electron Microscopy. The fluorescence was preserved in the resin. The block was imaged with a confocal microscope to identify the area of interest and to drive the FIB-SEM tomography acquisition. In this case, we targeted the intestinal region of an L1 larva of *C. elegans* whose cells expressed nuclear GFP. A) Overview of the resin block in the SEM. B) Overlay between the light and electron image (the field of view is indicated in the red square in A. C) Higher magnification of the ROI. This area is marked in panel B in yellow. D) The trench to expose the ROI was milled directly against the location of interest (arrow). E) Zoom in into the fluorescence microscopy maximum intensity profile around the targeted worm, in green the target intestinal cells, in red the neurons (white arrow). The white arrowheads highlight the dendrites. The green arrow is pointing at the intestinal cells which were targeted for FIB-SEM tomography. F) Overlay between the rendering of the confocal stack and the FIB-SEM tomogram.

In conclusion, we demonstrate that in contrast to all ions tested, oxygen represents a superior beam for milling the most currently used resins in the life sciences. In addition, the O-PFIB-SEM provides superior results in terms of data acquisition efficiency and image quality. Finally, we show that O-PFIB can be used to readily yield FIB-SEM tomograms in the context of CLEM experiments – opening up previously unavailable possibilities in this field such as targeted high-resolution imaging in large samples.

## Acknowledgements and author contributions

We acknowledge Dr. Chad Rue and Dr Jeremy Graham (ThermoFisher) for fruitful discussions on the alternative plasma sources. SG and GG collected the data, DK, AH, RL and VO prepared the samples, AdM, SG conceived the experiments, AdM, SG, MoB, JW, RP analized the data and wrote the manuscript. OK prepared the illustrations and provided the image registration algorithms.

